# Augmenting Bacterial Similarity Measures Using a Graph-Based Genome Representation

**DOI:** 10.1101/2024.04.08.588602

**Authors:** Vivek Ramanan, Indra Neil Sarkar

**Affiliations:** Center of Computational Molecular Biology, Brown University, Providence, RI, 02906, USA; Center for Biomedical Informatics, Brown University, Providence, RI, 02906, USA; Rhode Island Quality Institute, Providence, RI, 02906, USA

## Abstract

Relationships between bacterial taxa are traditionally defined using 16S rRNA nucleotide similarity or average nucleotide identity. Improvements in sequencing technology provides additional pairwise information on genome sequences, which may provide valuable information on genomic relationships. Mapping orthologous gene locations between genome pairs, known as synteny, is typically implemented in the discovery of new species and has not been systematically applied to bacterial genomes. Using a dataset of 378 bacterial genomes, we developed and tested a new measure of synteny similarity between a pair of genomes, which was scaled onto 16S rRNA distance using covariance matrices. Based on the input gene functions used (i.e., core, antibiotic resistance, and virulence), we observed varying topological arrangements of bacterial relationship networks by applying (1) complete linkage hierarchical clustering and (2) KNN graph structures to syntenic-scaled 16S data. Our metric improved clustering quality comparatively to state-of-the-art ANI metrics while preserving clustering assignments for the highest similarity relationships. Our findings indicate that syntenic relationships provide more granular and interpretable relationships for within-genera taxa compared to pairwise similarity measures, particularly in functional contexts.

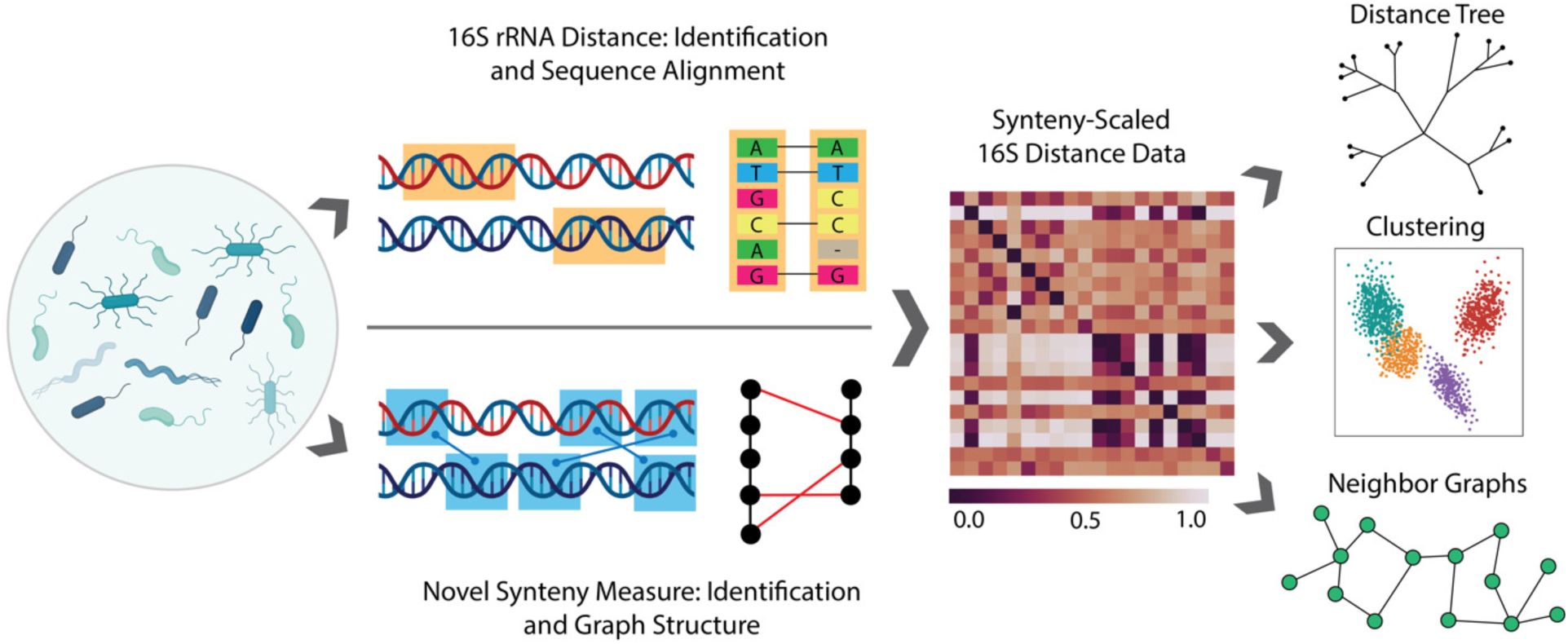
GRAPHICAL ABSTRACT. This method uses a novel measure based on shared synteny between bacterial genomes to scale 16S rRNA distance values. Created with BioRender.com.

## INTRODUCTION

16S ribosomal RNA regions are used to identify bacteria and form the foundation for phylogenetic relationships between bacterial groups [1, 2]. 16S rRNA analyses use variable regions of the 16S region to identify groups of similar sequences [3]. The level of identification (e.g., strain, species, genus) depends on sequencing power. However, improvements in technology expand the sequencing potential beyond the variable region of the 16S gene to the entire 16S region, as well as to entire genomes [4]. Studies have shown that 16S-based analysis is not infallible and does not always corroborate other forms of phylogenetics or taxonomy [5, 6]. 16S analyses rely on reference databases and heuristic clustering into ‘operational taxonomic units’ (OTUs), which can remove individual genomic sequences in favor of a consensus sequence [7]. While 16S regions continue to be the leading form of identification in bacteria, there are also numerous pairwise data that are often analyzed post 16S-identification or separate from 16S analysis. There is therefore potential to combine or analyze 16S alongside other data.

Matrix transformation is a mathematical approach that combines a pairwise matrix with another equally sized matrix. This approach transforms original data into a matrix that contains the variation of both input matrices enabling the combination to be analyzed as a single matrix. In the context of 16S, full genome information offers the potential of additional types of pairwise data. One example of the phenomenon of sequencing improvement is full-genome alignments, possible with tools such as MASH or FastANI [8, 9]. Here, entire genomes can be aligned in computationally efficient ways to gather similarity or distance scores between a pair of genomes. A limitation of this strategy is that distance scores are only provided under a certain threshold, which affects pairs that are not closely related. Contemporary metrics such as SKANI also include the usage of orthologous segment in average nucleotide identity (ANI) calculation, improving clustering quality of within-species phylogeny but is limited to calculating 82% ANI and higher [10].

Synteny is a comparative genomics approach that aligns shared segments of DNA between a pair of genomes, highlighting differences in a shared segment location [11]. The syntenic approach has been used as a visual representation for rearrangements, discovery of new shared segments, DNA order changes due to evolution, and genomic dynamics of subspecies [12–14]. A range of tools exist that provide synteny block construction (e.g., Sibelia) and visualization (e.g., Synteny Portal) [15, 16]. Synteny is used in targeted analyses and, to date, has not been employed in large scale analyses across a representative set of bacteria. Given synteny can provide a representation of similarity between two related genomes, we propose a two-fold approach for analyzing synteny: (1) characterize synteny as a representation for similarity between two related genomes, and (2) use syntenic data as an augmentation to 16S data using matrix transformation. Our synteny measure, which is based on a geometric graph structure, uses orthologous genes as the connection points between a given pair of genomes. We test the impact of this measure by transforming 16S rRNA data to demonstrate changes in clustering and graphical results, which can be used to understand new relationships between bacteria based on different data contexts. We compare this with state-of-the-art techniques with ANI calculation to test the difference in both clustering quality and clustering results.

## MATERIALS AND METHODS

### Data Acquisition from GenBank and Ortholog Construction

Bacterial genomes from GenBank were downloaded using ncbi-genome-download [17]. For each bacterial species, one strain was chosen at random, after which only genomes with 16S genes were chosen. CheckM was used to evaluate genome completeness and contamination, of which genomes with completeness > 90% and contamination < 5% were used for the study [18]. This resulted in 378 bacterial genomes in total for analysis. Genomes were organized in both GenBank Flat File and FASTA formats for further analysis. Taxonomic data on each species was gathered using NCBI Taxonomy.

### Core Gene Ortholog Construction

Core genes were identified using the UBCG2 dataset [19]. Core gene names and functions were taken from UBCG2 and compared with annotated genes and gene functions in GenBank flat files to identify core genes from the database. Identified genes were BLASTed against the entire gene database to identify orthologs, of which orthologs with greater than 95% nucleotide identity and a base pair length between 500 and 2500 base pairs were kept.

### Comparative Distance Score Calculation: 16S, MASH, and ANI

16S rRNA genes were identified from GenBank annotated flat files, which have been computationally predicted using protein homology. These genes were compared to the SILVA ribosomal RNA gene database project to confirm 16S identity [20]. For each pair of genomes, the MASH distance between each set of 16S genes per gene was calculated [8]. Whole genome MASH distances were also calculated using FASTA versions of sample genomes and ANI values calculated using SKANI on FASTA files. MASH distances were organized into a pairwise distance matrix, where the column and row indicated the species pair. The MASH distance was also validated against known taxonomic distance based on species name.

### Synteny Graph Structure and Measure

The construction of the pair synteny graph requires (1) a relative ordered set of genes for each of the genomes and (2) a set of at least 2 orthologous gene pairs between the two genomes. First, two linear graphs are created for each genome, after which edges are drawn between the orthologous gene pairs. Next, the cosine similarity of every combination pair of orthologs is calculated, using the relative order position of gene in genome A and the gene in the genome B (Figure 1). The average of the array of cosine similarities is calculated as the synteny similarity for that arrangement. To perform the other rearrangements to simulate circularity of genomes, a single pair is chosen as the pivot and arranged for each of the pair genes to be the top gene of the linear genome graph. The other genes are respectively ordered underneath the pivot gene of the pivot pair. The cosine similarities of every pair of orthologs are calculated and the average cosine similarity is produced. To find the final synteny similarity, the median synteny similarity across all rearrangements is used. Median synteny similarity for each pair of genomes was organized into a pairwise similarity matrix, of which the distance matrix was calculated by subtracting every matrix value from 1. Only core genes orthologs were used in the first iteration of the synteny similarity matrix, after which other functional gene group orthologs were used.

**Figure 1.**
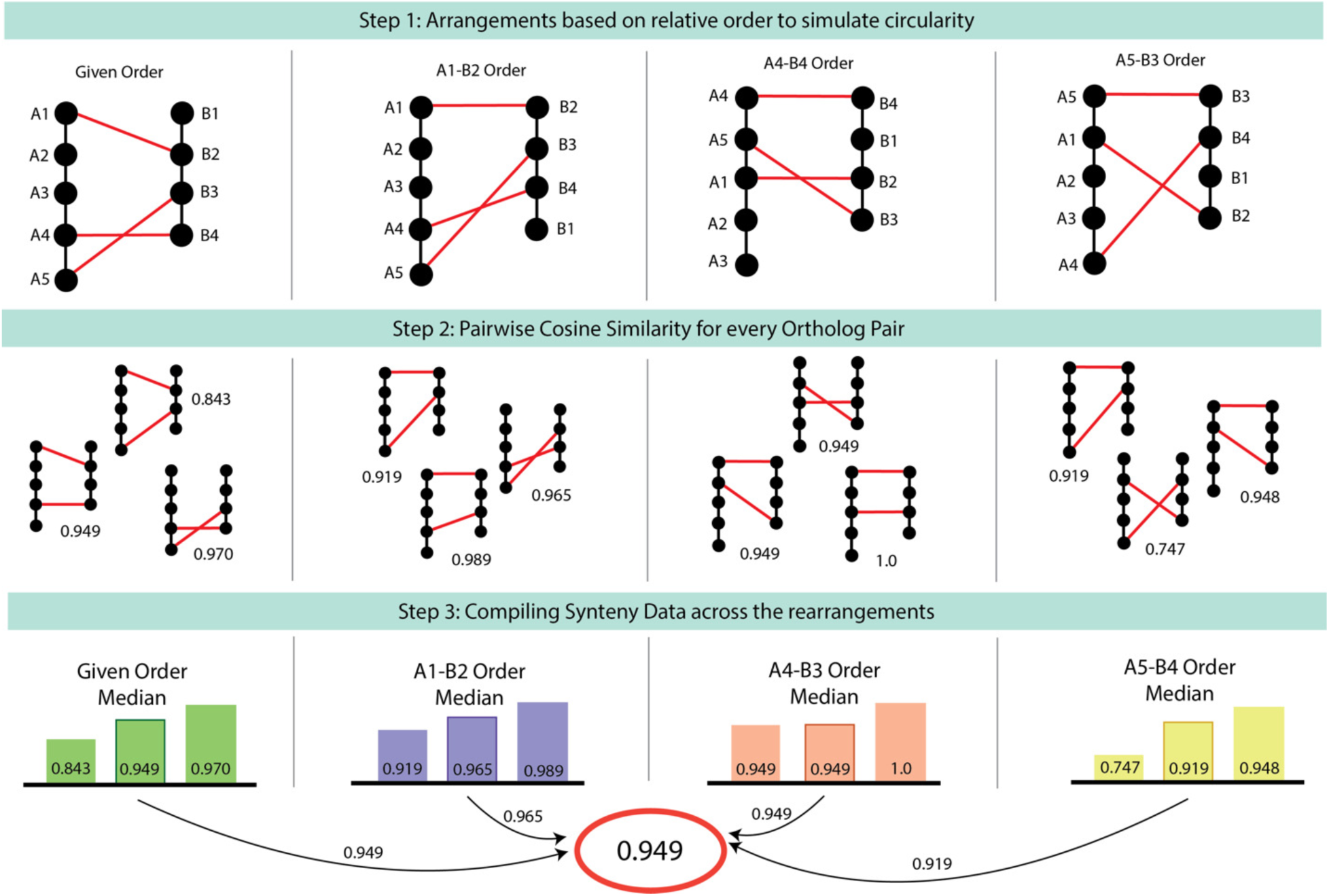
Visualization of the synteny similarity process. In Step 1, multiple arrangements are created based on the ortholog set given, where the first order is the given order from the GenBank flat files and each following order is based on a chosen ortholog as the pivot. In Step 2, the cosine similarity for each pair of orthologs is calculated, where values closer to 1 indicate similar positions of the orthologs and values closer to 0 indicate lower similarity. Finally, in step 3, the values are compiled per arrangement to find the median, then the median amongst all the arrangements are chosen. Median values were used to reduce skew based on a wide range of cosine similarity values.

### Synteny Coverage Metric

Synteny coverage was calculated as an asymmetric pairwise measure, equaling the summed length of the shared nucleotide blocks of a pair divided by the total genome length of the chosen genome [21]. Therefore, this measure is different for each of the genomes used in the pairwise measure.

### Random Forest Prediction

Random forest models were trained on a combination of 16S, whole genome distance, or syntenic distance (with all combinatoric possibilities) to predict taxonomic distance. Taxonomic distance was formed as a binary variable for each Linnean taxonomic level (i.e., kingdom, phylum, class, order, family, genus) and an individual model was trained for each level. Accuracy and ROC curves were calculated for every model possibility.

### Augmenting 16S data with Syntenic Similarity

To perform augmentation of the 16S MASH distance matrix, we used the covariance matrix of the synteny distance matrix. The dot product of the 16S MASH distance matrix and the covariance matrix of the synteny distance matrix resulted in the augmented 16S synteny distance matrix, which was normalized to range between 0 and 1. 0 indicated complete similarity based on 16S and synteny data, while 1 indicated no similarity. The completed matrix is symmetric, in which the diagonals indicate complete similarity for synteny of the same genome.

### Hierarchical Clustering

Complete linkage hierarchical clustering was performed on the synteny-scaled 16S distance matrix, the original 16S distance matrix, and the ANI distance matrix. Dendrograms were visualized using unrooted trees in ggTree and labeled using the bacterial taxonomic phyla [22]. The cluster cutoff number was varied to analyze metrics over a single experimental variable. To compare the results of the clustering groups, the Rand score and silhouette scores were all calculated. Rand scores form a similarity score for two clusterings when the matches between clusters are not known, by accounting for all combinations of cluster pairs between the two. Rand scores range from 0 to 1. Silhouette scores calculate the quality of clusters and range from - 1 to 1, with a value of 1 indicating a high-quality cluster. Metrics were compared across 16S data alone, ANI data, and the novel synteny metric. In addition, due to sparsity of data in ANI, two other reduced data versions of the synteny metric were used. One contained only values greater than 82% to match the threshold value of ANI and the other only contained the values for non-zero pairs in the ANI matrix. These were termed “Thr” and “Rem” respectively. The comparison was performed using ANI and synteny coverage in place of the synteny metric, where it was scaled using the covariance metric schema against 16S to calculate silhouette score.

### KNN Graphs

K-Nearest Neighbor (KNN) graphs were constructed for the original 16S distance matrix, the synteny-scaled 16S distance matrix, and the ANI matrix using NetworkX. Using the ‘distance’ mode, the normalized KNN array was visualized as a graph using the Kamada Kawai Layout of NetworkX. Visualized graphs were labeled with the taxonomic phyla or the taxonomic class. Network quality was calculated using modularity. The networks were compared using (1) Jaccard Similarity, (2) Weighted Edge Jaccard Similarity, (3) Dice Coefficients, and (4) DeltaCon Distance [23]. The inverse value of the DeltaCon Distance was used to provide similarity. Additionally, DeltaCon node and edge attribution was performed [24]. The cluster cutoff was varied to visualize which cutoffs had highest similarity scores and modularity changes in quality. Community detection of the KNN graphs was performed as a parallel for hierarchical clustering with the Girvan-Newman algorithm, with uses iterative removal of edges based on shortest path [25]. Girvan-Newman communities were displayed as a dendrogram, where the number of communities are chosen by the algorithm.

### Functional Application

Four cohorts were chosen to apply the synteny measure and augment 16S data with. The gene functions of (1) Mobile Genetic Elements, (2) Virulence Factors, (3) Antibiotic Resistance, and (4) Metabolic Genes (consisting of short chain fatty acids and neurotransmitter genes), were sourced using a combination of pre-existing descriptions in GenBank Flat Files and separate databases. Mobile genetic elements were identified using a set of keywords (transposase, transposon, conjugative, integrase, integron, recombinase, conjugal, mobilization, recombination, and plasmid) from flat file descriptions. Antibiotic resistance nucleotide sequences were gathered from the CARD database and compared to database sequences using BLAST, of which sequences with identity above 90% [26]. The retrieved sequences were BLASTed against the entire database again to identify orthologues, using greater than 95% sequence similarity and a length between 500 and 2500 base pairs. Virulence factors went through the same process using the Virulence Factor Database (VFDB) [27]. The metabolic genes cohort was identified using a combination of gut-brain modules from *Vieira-Silva, et al.* and bile-acid metabolism from *Funabashi, et al.*, which were sourced as KEGG modules [28, 29]. Genes in the KEGG modules that matched the dataset’s species were downloaded and then BLAST’ed against the gene database to orthologs once again [30]. Each cohort of orthologs was filtered the same way as the original core gene cohort and synteny similarity was calculated for pairs of genomes that had at least two ortholog pairs. Random forest models were trained on functional synteny to predict taxonomy. The same augmentation process was used to form 16S synteny-scaled distance matrices for each functional cohort, with only values above the 82% threshold. Hierarchical clustering was performed, with silhouette scores and Rand scores calculated. KNN graphs were also visualized with weighted Jaccard and DeltaCon coefficients calculated between the original 16S and original 16S synteny-scaled matrix.

## RESULTS

### GenBank public data provided genomic, 16S, and core gene data

Our analysis focused on 378 genomes retrieved from GenBank, which represented ten phyla (*Actinomycetota, Bacteroidota, Campylobacterota, Chlamydiota, Bacillota, Fusobacteriota, Pseudomonadota, Spirochaeota, Mycoplasmatota,* and *Verrucomicrobiota*), 19 classes, 125 genera, and 378 species. *Pseudomonadota* and *Bacillota* had the highest representation (with 135 genomes each), followed by *Actinomycetota* (n=42) and *Bacteroidota* (n=29). The other phyla had under 15 genomes per phyla, with *Verrucomicrobiota* having only a single genome. Using the 16S sequences from each genome, we calculated the MASH distance between every pair of genomes, which accounted for multiple unique 16S genes per genome [31, 32]. Core genes were identified using the UBCG2 database, resulting in 34,051,278 genes across the database, roughly averaging to 70 genes per genome.

### Synteny similarity measure indicates genus-level dynamics among bacteria

The developed syntenic measure ranged from 0 to 1, where 1 indicated complete similarity and 0 indicated complete dissimilarity between a given genome pair. The proposed method involves forming multiple rearrangements of the relative gene order, followed by pairwise cosine similarity values for each ortholog pair, and a compilation of data values across the pairs and arrangements (Figure 1). In the dataset used for this study, synteny values for core genes were found for pairs within the same phyla, representing the phyla of *Bacillota*, *Actinomycetota*, *Pseudomonadota*, *Bacteroidota*, and *Spirochaetota* (Figure 2A). Smaller sub-networks were identified from the greater network of synteny, where each edge represented a synteny similarity and is weighted by the similarity. The two largest sub-networks were of *Pseudomonadota* and *Bacteroidota*, the largest comprised of the species in *Gammaproteobacteria* class and the second-largest formed of species in the *Bacteroidales* order (Supplementary Figure 6,7). *Pseudomonadota*, *Bacteroidota*, and *Bacillota* made up most of the data, with *Actinomycetota* and *Spirochaeta* making up smaller sets. The distribution of synteny similarity values was therefore separated into within-genus pairs and out-of-genus pairs. Median synteny similarity values differentiated between pairs in the same genus and pairs outside of the same genus (Figure 2B). The sparsity of the synteny distance matrix augments 16S distance by using a covariance matrix-based transformation (Figure 2C). The 16S and synteny distance matrices additionally had low correlation (Spearman = 0.149), indicating different informative values in either matrix. Out of 143,641 total pairwise combinations, there were 553 non-zero values in the sparse similarity matrix and 143,625 non-zero values in the 16S distance matrix. The final scaled matrix has 143,262 non-zero values (0.02% sparsity), increasing the original 16S matrix from 58% sparsity with the synteny matrix of 99% sparsity. The ANI matrix also had 99% sparsity as well, which indicated that sparse version of the scaled synteny metric were necessary for comparison. Two reduced cohorts were formed, labeled “Thr” and “Rem”. In Rem, data was removed to only contain the same position values as the ANI matrix (99% sparsity) and in Thr, data was only used at a threshold above 82% similar to ANI field metrics (77% sparsity). Random forest models, which are apt for identifying underlying nonlinear trends, were trained to predict taxonomic distance for data associated with either 16S distance or synteny distance (1 – synteny similarity). In all six taxonomic levels, 16S distance performed the best consistently, which lowered at higher taxonomic levels (e.g., phyla, class). For the synteny similarity, only the prediction for the genus level was greater than random, while all others resembled random chance (16S AUC: 0.7229, Synteny: 0.649, Combined: 0.749) (Figure 2D).

**Figure 2.**
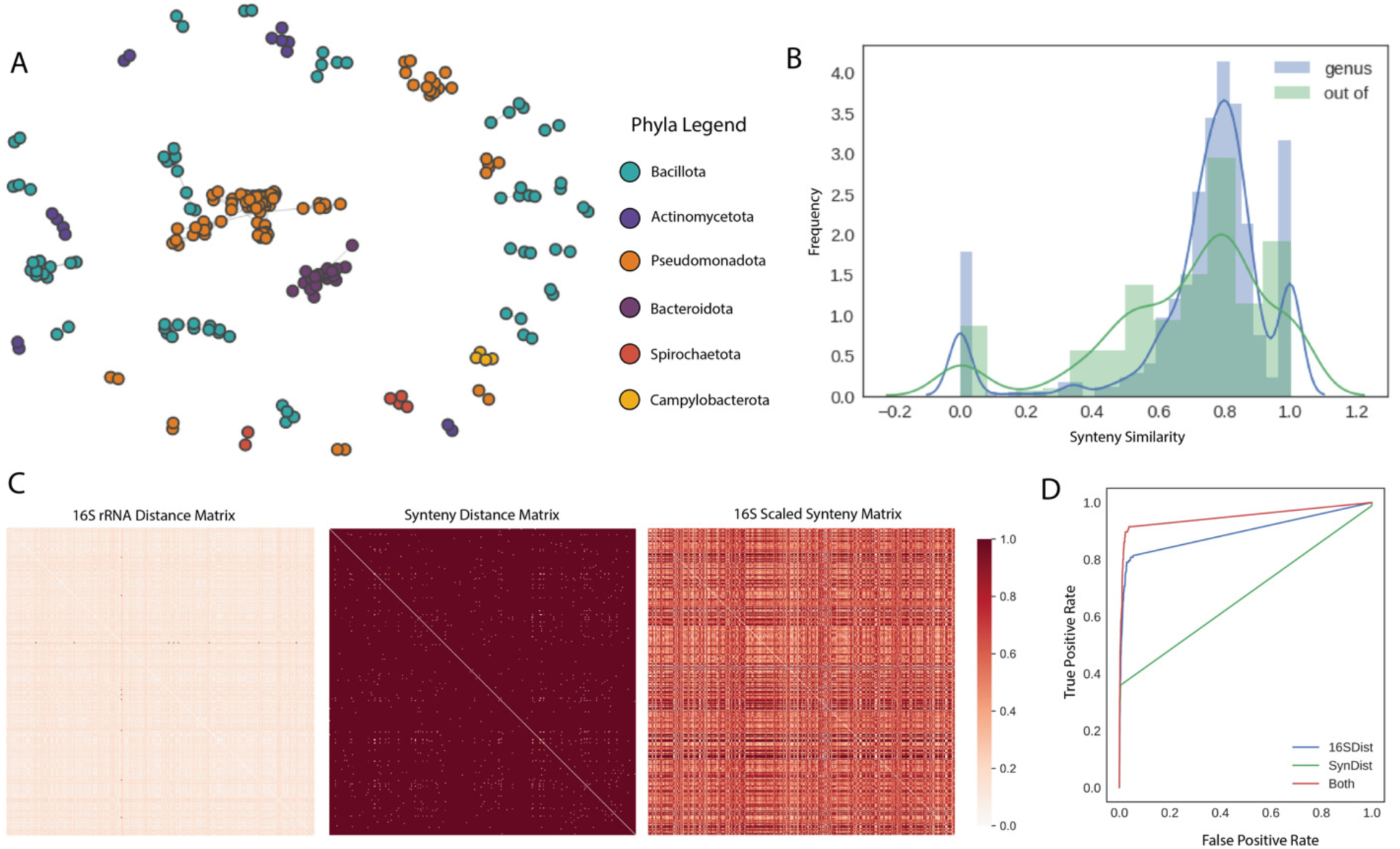
(A) A network diagram showing organization of synteny similarity measure, where nodes represent individual genomes of species and labeled by taxonomic phyla. (B) The count histogram of the synteny similarity measure differentiates between pairs that share the same genus versus pairs that have more taxonomic separation. (C) Each heatmap depicts a particular step of the data analysis, the first being the original 16S distance matrix followed by the synteny distance matrix (calculated as 1 3 synteny similarity). The last heatmap displayed the final normalized matrix, the combination of the two previous distance matrices. (D) The individual ROC curves for the random forest model trained on taxonomic genus (0 = pair in the same genus, 1 = pair in the same genus). Each curve represents the data input given (16SDist: 16S MASH distance, SynDist: synteny distance, Both: 16S and synteny distance values).

### Hierarchical clustering of 16S Scaled-Synteny data displays shifting of clusters

Using the 16S scaled-synteny, original 16S, and ANI distance matrices, we performed complete linkage hierarchical clustering. Silhouette scores were used to visualize clustering quality, ranging from +1 to −1 with −1 indicating poor clustering quality and +1 indicating high clustering quality. When varying the cluster cutoff number, the novel synteny metric performed the most consistently for cluster quality via silhouette score in the order of thr (Thresholded metric), Rem (removed metric), and the novel synteny metric termed “Syn” (Figure 3C). The synteny metric alone without the 16S data performed similarly to the ANI metric as well. The scaling of 16S with ANI data in the same covariance format resulted in negative silhouette scores, indicating reduced clustering assignments (Supplementary Figure 5). Synteny coverage also performed negatively compared to 16S and the synteny metric. Resulting dendrograms used cluster cutoff of 5, where the highest consistent silhouette score was observed, showed differentiated clustering between the scaled and original distance matrices (Figure 3A and 3B). The dendrogram from the original matrix clustered *Pseudomonadota* and *Bacteroidota*, while *Actinomycetota* and *Bacillota* were clustered into the same general grouping but remained distinct (Figure 3A). The dendrogram from the scaled matrix slightly shifted the positions of some groupings (Figure 3B). Most *Pseudomonadota* remained separate, while a small group clustered with *Bacteroidota, Spirochaetota,* and *Chlamydiae* interestingly. The combined clustering of *Bacillota* and *Actinomycetota* remained, with a smaller group cluster further away near *Mycoplasmatota*. *Bacteroidota* also clustered further away. In both dendrograms, *Verrucomicrobiota* clustered the furthest from all other species and remained an outgroup. The remaining data groups of ANI, Thr, and Rem were extremely sparse, resulting in incomplete dendrograms.

**Figure 3.**
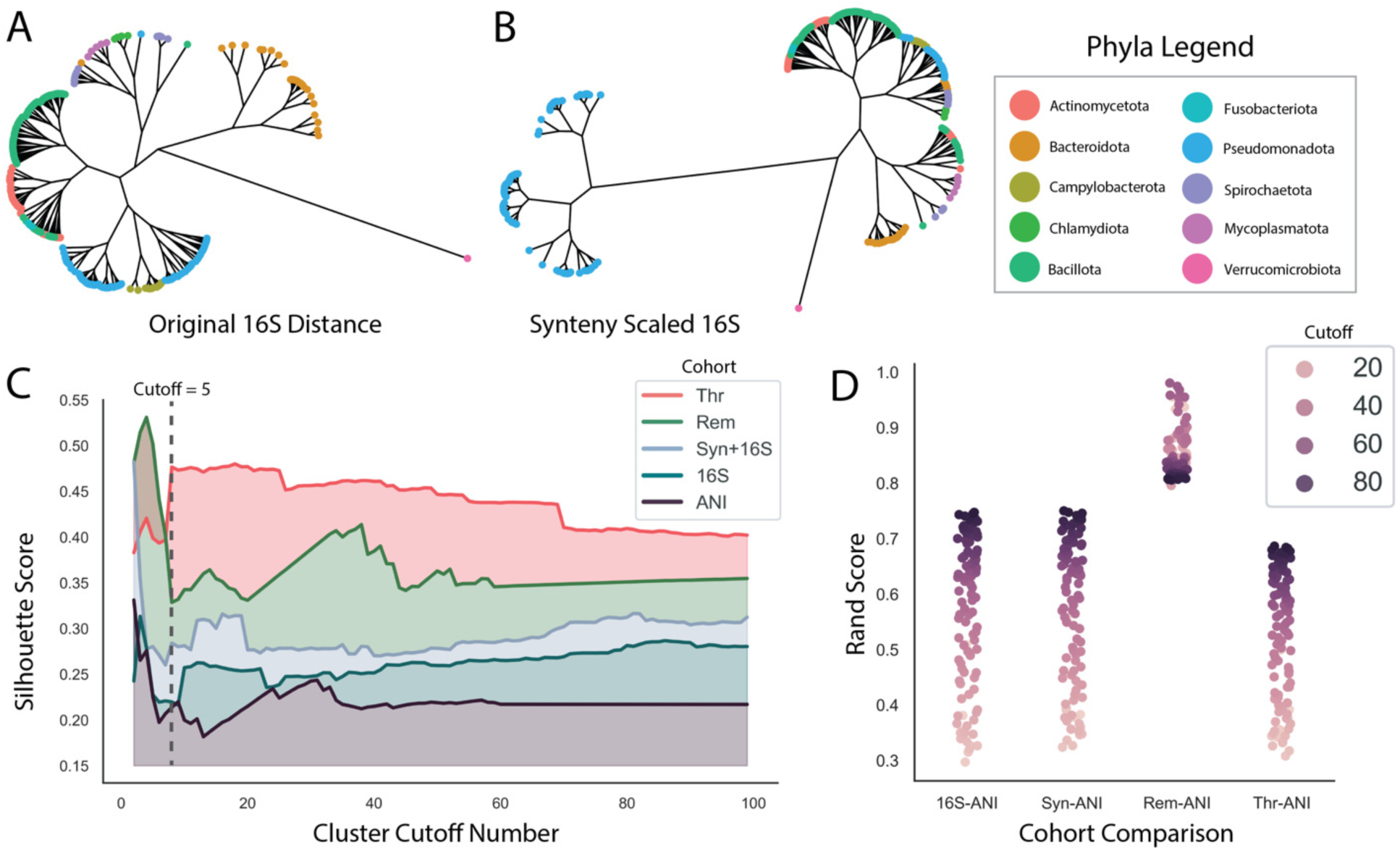
Hierarchical clustering displays increased clustering quality with synteny metric compared to ANI. (A, B) The results of hierarchical clustering on the data are represented as an unrooted dendrogram at cluster cutoff of 5, with the taxonomic phyla labeled. A represents the original 16S data and B represents the synteny scaled 16S data. (C) Varying the cluster cutoff number from 1 to 100 with the silhouette score across the 5 different versions of comparisons (ANI: average nucleotide identity, Syn + 16S: the synteny scaled 16S data, Thr: the Syn+16S cohort with values only greater than 82% threshold, Rem: the Syn+16S data with the same positional values as the ANI data). (D) Comparisons between each data group and the ANI data, defining the Rand score across the same cluster cutoff modulation. The Rand score shows the similarity of clustering while accounting for possible different cluster numberings. The hue of the point represents the cluster cutoff number.

In addition, comparisons between clustering results were performed against ANI as a ground truth standard (Figure 3D). The rand score measures similarity of clustering decisions, in which all possible combination of cluster labels between two sets are considered to account for differential cluster number and labeling. Scores ranged from 0 to 1, with 1 indicating total similarity and 0 indicating no similarity. The highest rand score is in the Rem-ANI comparison, reflecting the fact that the Rem matrix contains the same position values as the ANI matrix. The 16S and ANI comparison showed the next highest Rand, with the Thr and ANI comparison having a slightly lower Rand score.

### KNN Graphs offer an alternate form of clustering and visualization of synteny scaling

A secondary approach to analyze differences between the original, synteny-scaled, and ANI data was through K-nearest neighbor (KNN) graphs. KNN graphs were generated for all distance matrices (16S, SYN, REM, THR, and ANI) across k ranging to 15 (Figure 4D), where a midpoint of k = 10 was used for visualization. Nodes indicated species and edges were weighted by the k neighbors algorithm based on the distance values given. Labels were chosen based on taxonomic phyla and class. Visual differences between two networks were based additionally on layout structure, using the Kamada-Kawai path-length cost function. The phyla labeled networks show a distinct set of groupings based on phyla (Supplementary Figure 8). Similar groups from the hierarchical clustering dendrograms were seen in these networks. In the original 16S network, *Pseudomonadota* separated into two separate groups, whereas in the scaled network, *Pseudomonadota* expanded out linearly (Figure 4A and 4B). When the taxonomic labeling was changed to class, this expansion was clearly based on *Gammaproteobacteria*. Many of the other groups remain consistent such as *Bacteroidota, Verrucomicrobiota,* and *Bacillota,* with the *Bacillota* composition splitting into *Bacilli* and *Clostridia* distinctly in both the 16S and scaled KNN graphs. We examined these differences further using the Weighted Jaccard and DeltaCon comparative frameworks. the DeltaCon similarity measure (0.928), which also assigns attribution to nodes and edges for the impact of difference between the two networks (Figure 4C). This edge and node attribution, when accumulated per taxonomic class and normalized, resulted in highest attribution to *Gammaproteobacteria*, *Bacilli*, *Clostridia*, *Actinomycetia*, and *Spirochaetia*. The lowest values of attribution were seen in *Verrucomicrobiae, Tissierelia, Erysipelotrichia,* and *Deltaproteobacteria.* In contrast, the KNN graph of the whole genome scaled 16S data shows much less consistency of taxonomic groupings other than *Bacillota* and *Pseudomonadota*, where in the rest of the groups are interspersed. Weighted Jaccard was used for speed to identify similarity between an individual network and the ANI network across the k value modulation (Figure 4E). Between the 4 groupings, the 16S-ANI has the highest similarity overall, while the Thr-ANI similarity starts with high similarity at low k values then decreases lower.

**Figure 4.**
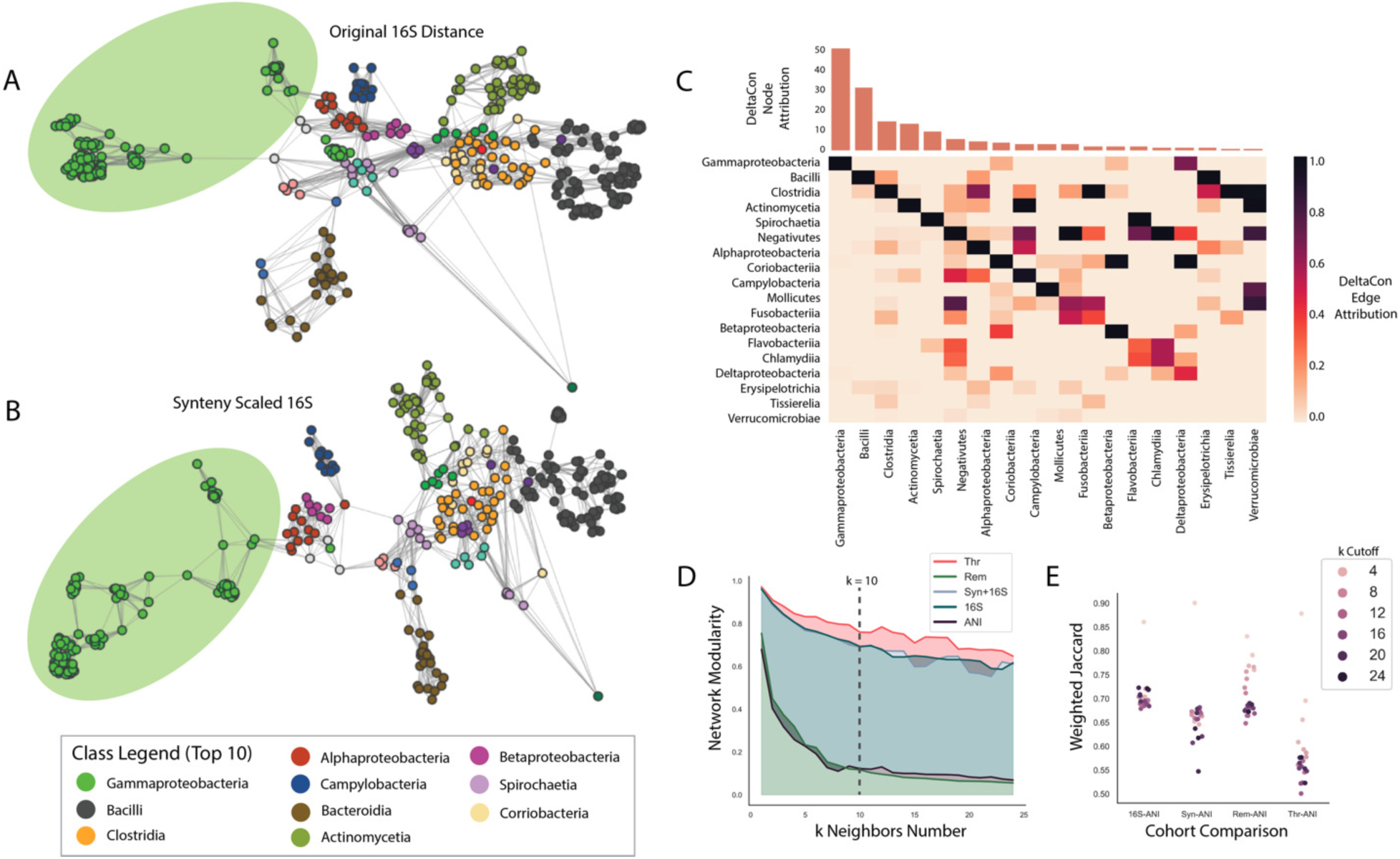
KNN graphs display connection differences of synteny-scaled data. Visualizations of KNN graphs on 16S and synteny-scaled 16S data are displayed in A and B, labeled based on taxonomic class. (A) represents the original 16S data while (B) displays synteny-scaled 16S data, where only the top 10 classes are displayed in the legend for brevity. The *Gammaproteobacteria* class is highlighted for relevance. (C) contains results of the DeltaCon network comparison method, where nodes and edges are given attribution to quanity their impact on the difference between two networks. Attribution values were accumulated into taxonomic class and then normalized. Node attribution is seen in the top bar graph, ordered based on frequence. Edge attribution is represented by a pairwise heatmap, where higher values indicate those pairwise edges held more importance to the difference between networks. (D) displays the network modularity over the increasing k variable for the KNN graphs. (E) shows the comparisons of paired networks against ANI networks using the weighted Jaccard score.

### Applying Syntenic Measure to Functional Gene Groups

Four functional gene cohorts were used to test the synteny measure and clustering frameworks, which were antibiotic resistance (ARG), mobile genetic elements (MGE), metabolic genes (MET), and virulence factors (VIR). The number of genes found per each genome represents the ubiquity or the specialization of these genes per bacterial group, in which antibiotic resistance and virulence factors tend to be more specific to specific bacterial groups, while metabolic genes and core genes tend to be more universal in terms of the numbers of genes present (Figure 5A). The distribution of synteny similarity measures was quite similar across all five functional groups, of which the original metric based on core genes has the lowest average, while the other four groups have similar averages but different ranges in terms of 1^st^ and 3^rd^ quartiles (Figure 5B). The same analyses were performed on the larger cohorts to identify differences between the original synteny metric based on core genes to the functional cohorts in terms of clustering quality and comparison to ANI. Hierarchical clustering modulations showed the highest performance with the ARG group, followed by MET, VIR, MGE, and finally the synteny metric (Figure 5C). Weighted Jaccard comparisons with the ANI network showed highest similarity between the MET-ANI, followed by VIR, MGE, and finally ARG (Figure 5D).

**Figure 5.**
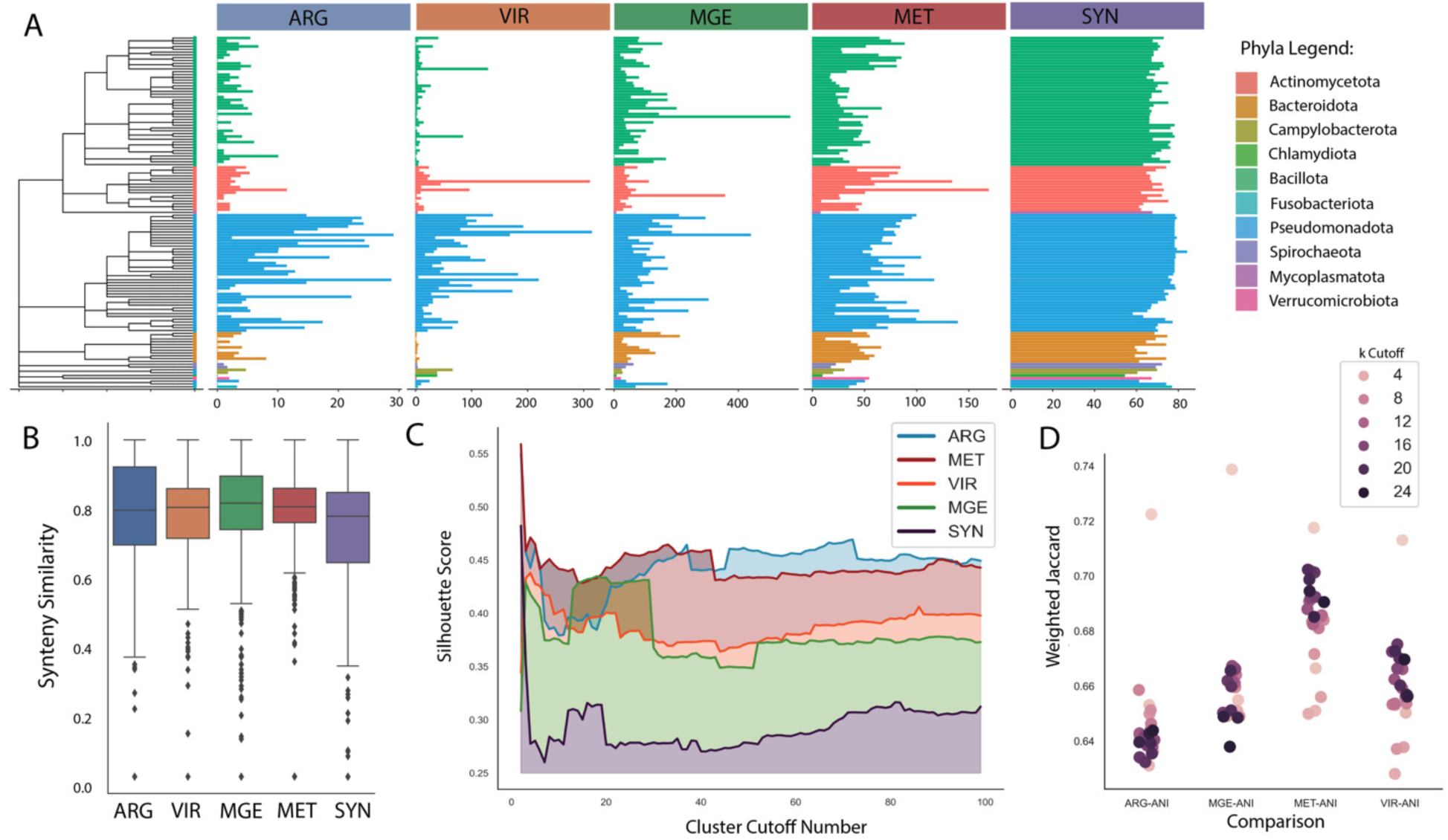
An overall representation of the results of the synteny scaling process on multiple functional gene cohorts. (A) Mean quantities of cohort genes found per genus, where ARG: Antibiotic Resistance, VIR: Virulence Factors, MGE: Mobile Genetic Elements, MET: Metabolic Genes, and COR: Core Genes. Color labels indicate taxonomic phyla. (B) Synteny similarity distributions as box plots for each functional cohort. (C) Results of hierarchical clustering across cluster cutoff increasing, indicating changes in clustering quality using silhouette scores. (D) Comparisons of cohort KNN graphs against the ANI network using weighted Jaccard, where higher values indicated greater similarity between two networks.

## DISCUSSION

The goal of this study was two-fold, to: (1) suggest an augmentation procedure for 16S rRNA data with a genomic pairwise measure, and (2) provide and test a novel measure based on genomic synteny. Two clustering procedures were used to compare differences in the original and transformed data, one using hierarchical clustering and other using k-nearest neighbor graphs. The ground truth of these clustering approaches ultimately is unknown, particularly given that historical bacterial taxonomic nomenclature has been based on limited 16S rRNA values and phenotypic data [33, 34]. Therefore, ANI was used as a proxy for ground truth given it is a field standard and consistent against genome fragmentation, to understand how individual versions of the metric fare against ANI.

The results of the synteny similarity measure depict dynamics that match taxonomic relationships. At the genus level, more synteny similarity values are seen within-genus as opposed to out-of-genus (Figure 2B). Based on the input data for this task (shared core genes orthologs between two genomes), this result validates the assumption that closer related genomes are more likely to share more genes. Therefore, there are more non-zero similarity values present for genomes that are in the same genus as opposed to those outside of the same genus. This also validates the random forest result that prediction only on the genus level is above random. The synteny similarity network also demonstrates that all synteny measures are within-phyla, indicating non-zero values for genome pairs that are more closely related than out-of-phyla pairs (Figure 2A). The sparsity of the final synteny matrix displays its value when mathematically combined to the original 16S matrix, creating a pairwise matrix that has more variation than the two input matrices (Figure 2C). The synteny measure on its own therefore does not indicate taxonomy other than being within-phyla and genera specific but is not necessarily predictive of these attributes. Instead, it is most valuable when scaled on other non-sparse data and significantly reduces matrix sparsity.

The hierarchical clustering and KNN graph approaches reveal different pieces of information with respect to the original 16S distance data and the ANI data. In the clustering approach, the quality of the clustering, determined by the silhouette score, is better for the synteny matrices in comparison to the ANI and original 16S matrix. This likely reflects the improvement in sparsity of the matrix values. In contrast, using the ANI matrix as an augmentation on 16S resulted in negative silhouette scores but similar rand scores as 16S and the synteny metric. The rand scores depict that higher cluster cutoff values lead to better rand scores, or similarity in clusters between ANI and the metric. Interestingly, the opposite occurs in the KNN graphs where lower k values result in higher similarities to the ANI network via the weighted Jaccard metric. Additionally, the Rem matrix performs similarly to ANI likely due to high sparsity of data as well. Therefore, between the sparsity of the Rem matrix and the lower similarities between ANI and the Thr matrix, the original synteny matrix represented the best combination of clustering quality and similarity to ANI for the functional cohorts. These results also highlight an interesting dynamic between the two clustering approaches. Hierarchical clustering draws out general dynamics across the entire dataset, particularly stabilizing and improving ANI similarity at higher cluster cutoff values. Whereas the KNN approach performs better at low K values, highlighting the strongest pairs. The choice of respective hyperparameter (k in KNN and the number of clusters in clustering) determines the interpretation potential and the type of information available in each complementary approach.

The application of these approaches to the functional cohort reveal that the choice of input plays a strong role in the value and interpretation of this metric. While originally core genes were solely used for the metric in the original synteny metric, we observed that antibiotic resistance and metabolism-based genes performed quite well in terms of clustering quality via silhouette scores, with values closer to the Thr matrix in Figure 3C. In addition, the metabolism cohort also had the highest weighted Jaccard similarity to the ANI networks. While the metabolism cohort has higher amounts of data compared to the core genes, the mobile genetic elements cohort also has higher data quantities yet does not perform as well. Therefore, it is possible that there is some underlying signal where metabolic genes are providing more granular detail on microbial relationships. Previous studies have shown the importance of core metabolic genes as necessary to the minimal bacterial gene set or “pan genome”, which is possibly being reflected here [35]. However, antibiotic resistance genes also show high clustering quality despite having lower amounts of genes and lower similarity to the ANI networks. Whereas the virulence cohort also shows higher similarity to the ANI networks via weighted Jaccard, possibly indicating that the virulence genes represent distances closer to ANI, whereas antibiotic resistance has less similarity. In both approaches, *Gammaproteobacteria* make the highest impact in terms of number of genes per cohort as well as most amount of synteny data, which historically encompass many pathogens as well as having the higher representation of antibiotic resistance and virulence in our dataset (Figure 5A). This is unsurprising given that the data has the most amount of genomes for this group of bacteria, potentially due to the clinical bias of GenBank [36]. This is particularly visible in the change in KNN graph structure between the 16S data and synteny-scaled 16S data (Figure 4A and 4B). However, despite this bias, there are also an equivalent number of *Bacillota* genomes present in the data and consistent core genes that made up the original synteny metric, which do not significantly change structure. Thus, this phenomenon depicts that as data quality and number of genomes per bacterial class increases, visible changes in clustering and network structure truly represent changes in the genomic data, rather than changing solely as data increases. Metabolic genes are most successful across all groups at replicating ANI relationships and improving clustering quality. In contrast, antibiotic resistance genes can improve novel relationships between bacteria that share resistance, which are less likely to be found by ANI and 16S measures. Thus, it’s therefore possible to consider this tool in a gene function distribution context as well, to see how functionality affects the final similarity to ANI and whether those functions reflect if a pair of bacteria are known to be related or not.

Current taxonomy and identification improvements have focused on using whole genome alignments instead of 16S sequences, particularly for pathogen outbreak tracking and variant differentiation [37, 38]. Some techniques make use of solely vertically transferred genes, which illustrates a choice of genes for phylogenetics [39]. Computational methods have also been designed to combine sequencing data from different PCR-amplified 16S rRNA regions to increase resolution [40]. Methods in other species have made use of multiple rRNA sequences (18S, 16S, and 28S) along with cytochrome C to define phylogenetic relationships by combining the alignments of multiple conserved genes [41]. The synteny measure also differs from average nucleotide identity by providing detailed information for bacterial relationships that expand a single distance value. Few studies have explored functions as features of genomic relationships, but has been explored in the context of horizontal gene transfer [42]. As bacterial nomenclature also continues to change and develop with updated information, this augmentation method can provide context-specific relationships that are independent of nomenclature changes [43]. We are unaware of other methods that mathematically combine pairwise data with other forms of genomic data or characterize synteny as a pairwise similarity value.

Our proposed method comprises of an augmentation approach and a similarity measure that can be applied to pre-existing pairwise data. The choice of genes used for the similarity measure can play a role in determining the strength of relationships between already related bacteria. Some potential use-cases for this method can include: (1) finding relatives to a novel pathogen based on synteny of antibiotic resistance or virulence genes, (2) identifying bacteria with similar horizontal gene transfer profiles based on the syntenic similarity of transferred genes, and (3) finding the most functionally similar symbionts in different communities which share the same genes and syntenic attributes of those genes. As sequencing technology improves, many outstanding questions remain about how bacterial analysis should be conducted in the future. Is taxonomy relevant for clinical decision making [44]? Will whole genome sequencing replace 16S for bacterial identification in the future? How can reference-free approaches be considered instead of those that rely on genome references? We suggest that as bacterial genomic data grows in both quantity and quality, augmentation approaches can be applied to understand relationships in context-dependent environments. The proposed synteny similarity measure is one such example that makes use of available data, building on pre-existing knowledge of bacterial taxonomy and potentially be applied to functional and clinical contexts.

## CONCLUSION

In this study, we propose a data scaling method, which adds a novel similarity measure to traditional 16S rRNA distance scores, using matrix transformation. Our pairwise measure is based on a graphical structure of a bacterial genome, using ortholog location to form a numerical representation of synteny. Analyzing synteny-scaled 16S rRNA data in comparison to 16S rRNA and ANI data shows that our approach improves on clustering quality, while also retaining ANI relationships, particularly when using metabolic genes as the metric input. In contrast, antibiotic resistance genes can possibly unveil novel relationships that were not previously considered in clinical contexts as well. Ultimately, the choice of clustering method and input gene function determines the interpretation of relationship between bacteria, making this method a context-aware and dynamic approach that utilizes a novel genomic attribute to determine bacterial relationships.

## DATA AVAILABILITY

The data underlying this article were accessed from the GenBank Public Repository, hosted by the National Center of Biotechnology Information (NCBI). The datasets are available in the article and in its online supplementary material. Sequence data for each cohort will be shared on reasonable request to the corresponding author.

The code is publicly available at https://github.com/vivekramanan/synteny-scaling

## SUPPLEMENTARY DATA

Supplementary Data are available online.

## AUTHOR CONTRIBUTIONS

Vivek Ramanan: Conceptualization, Formal analysis, Methodology, Validation, Writing— original draft. Indra Neil Sarkar: Conceptualization, Methodology, Writing—review & editing.

## ACKNOWLEDGEMENTS

We thank Dr. Sorin Istrail for his support in developing the graphical structure for synteny similarity. We thank Drs. Tal Korem, Lorin Crawford, Shipra Vaishnava, Peter Belenky, and Rodrigo Bacigalupe for their inputs into the research.

## FUNDING

This project was supported in part by Institutional Development Award Number U54GM115677 from the National Institute of General Medical Sciences of the National Institutes of Health, which funds Advance Clinical and Translational Research (Advance-CTR), as well as the Predoctoral Training Program in Biological Data Science at Brown University from the National Institutes of Health (5T32GM128596-05). The content is solely the responsibility of the authors and does not necessarily represent the official views of the National Institutes of Health.

## CONFLICT OF INTEREST

No conflicts of interest to declare.

---

**Supplementary Figure 1.**
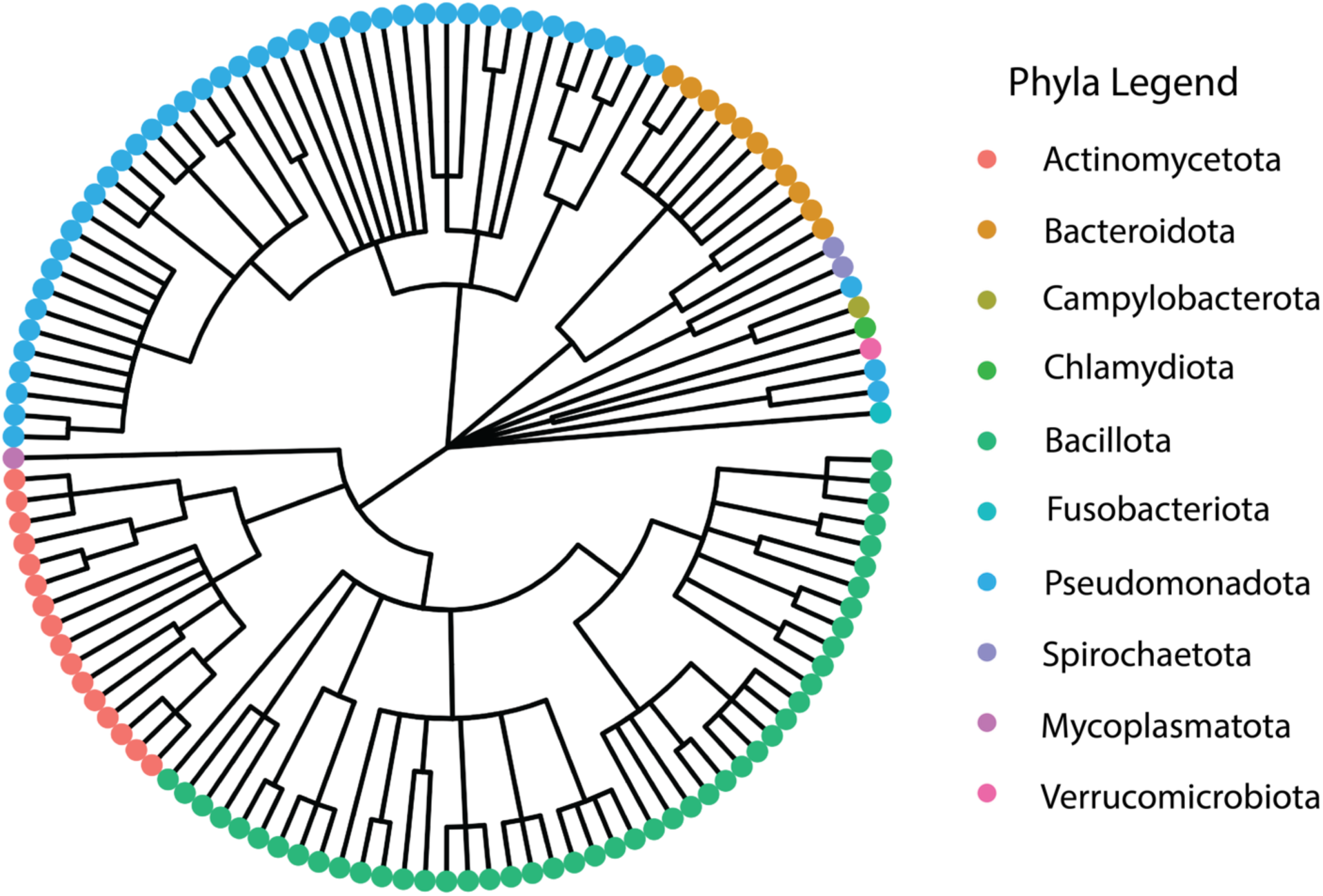
Composition of GenBank genome data, representing 10 phyla of bacteria. The largest groups comprise of *Pseudomonadota, Bacillota, Actinomycetota,* and *Bacteroidota.* The phylogenetic tree structure was created with NCBI Common Tree, with the genera used for input.

**Supplementary Figure 2.**
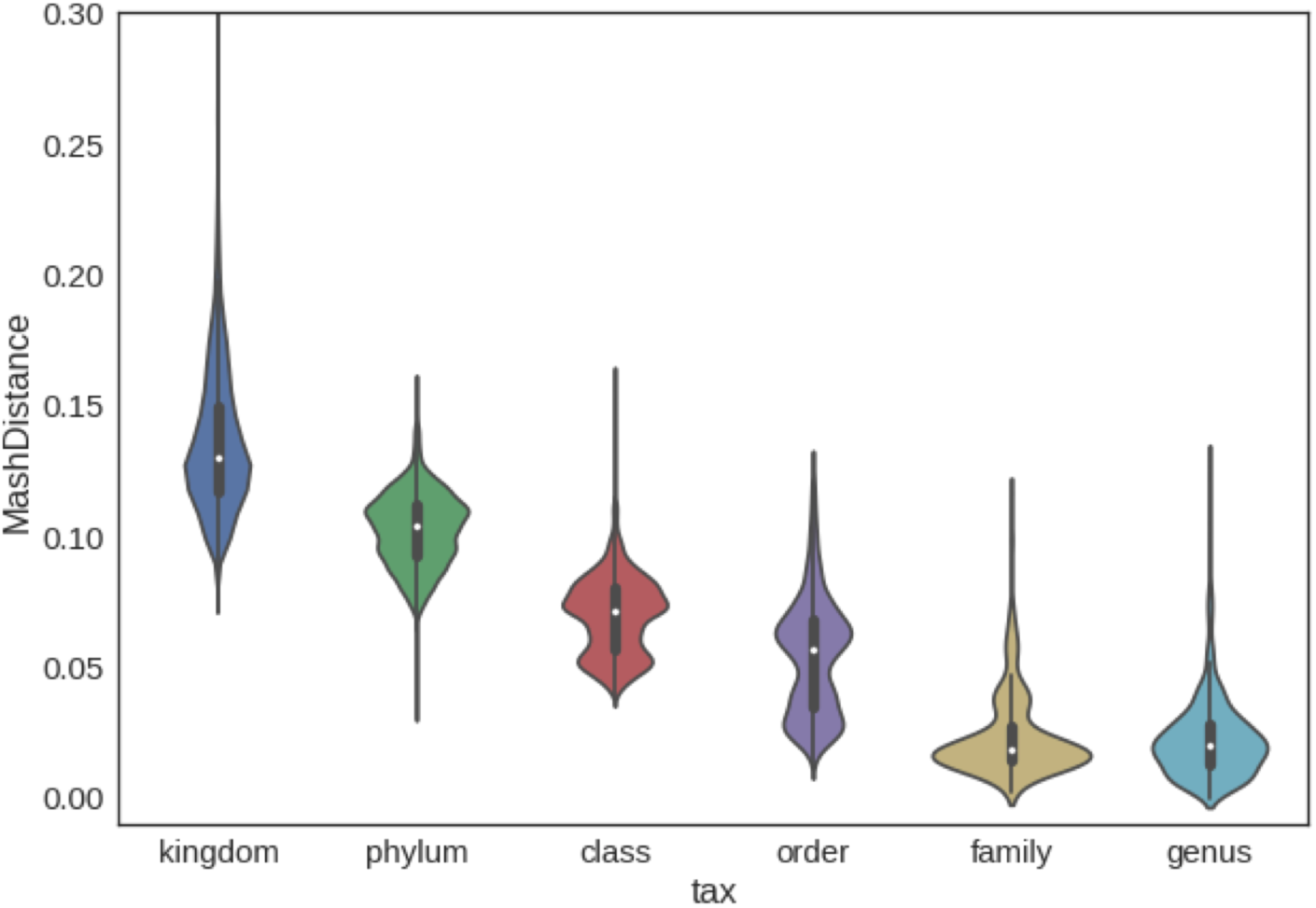
Distribution of 16S rRNA MASH distances, grouped by lowest shared taxonomic group between every pair of genomes (tax). For example, if two genomes are in two different phyla, the group would be labeled as kingdom. The lowest grouping possible is genus, where two genomes would be two different species within the same genus. Mash distances of 1 are not included in the plot, as these are rough calculations based on the MASH algorithm and do not represent exact distance.

**Supplementary Figure 3.**
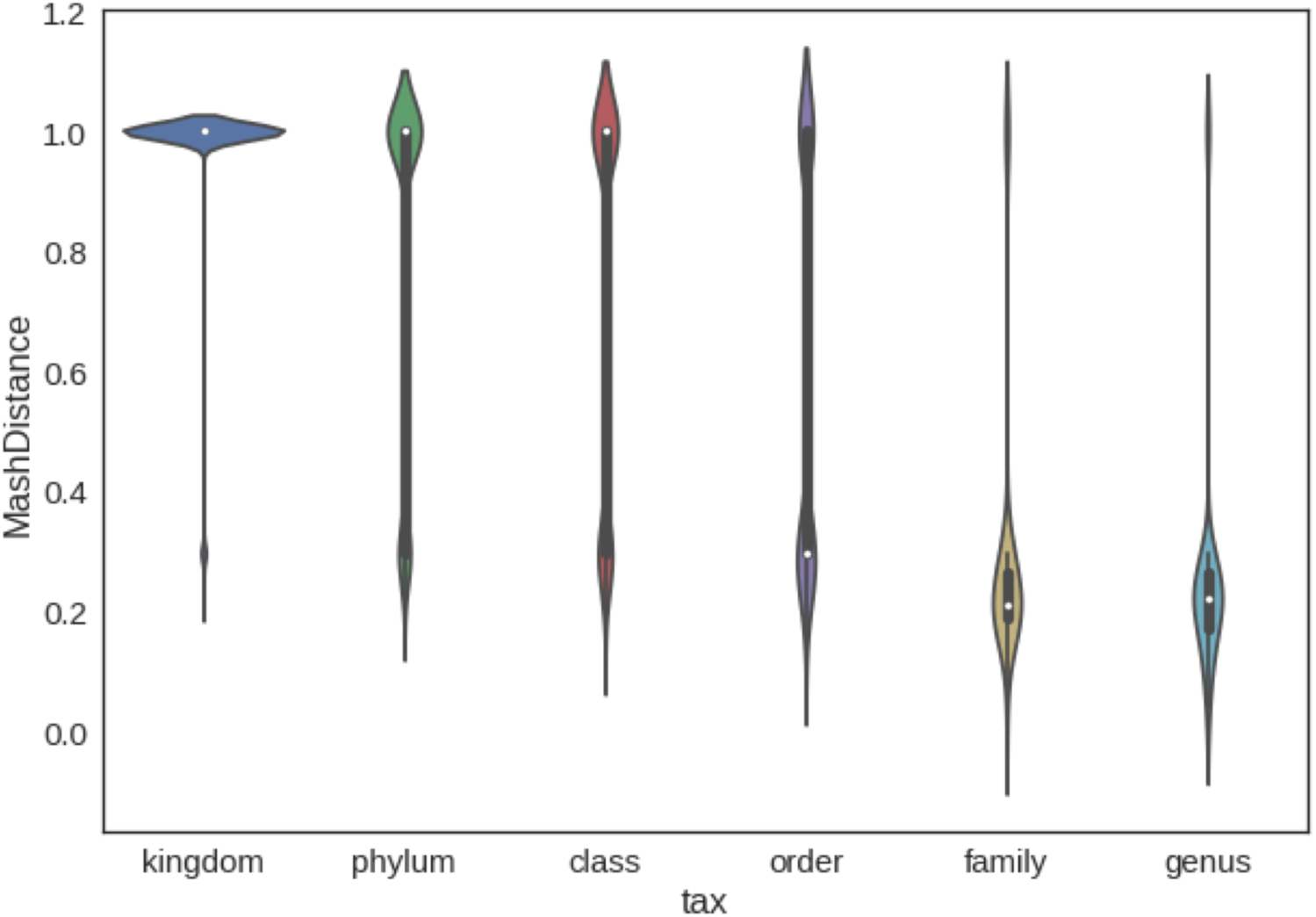
Distribution of whole genome MASH distances, grouped by lowest shared taxonomic group between every pair of genomes (tax). For example, if two genomes are in two different phyla, the group would be labeled as kingdom. The lowest grouping possible is genus, where two genomes would be two different species within the same genus.

**Supplementary Figure 4.**
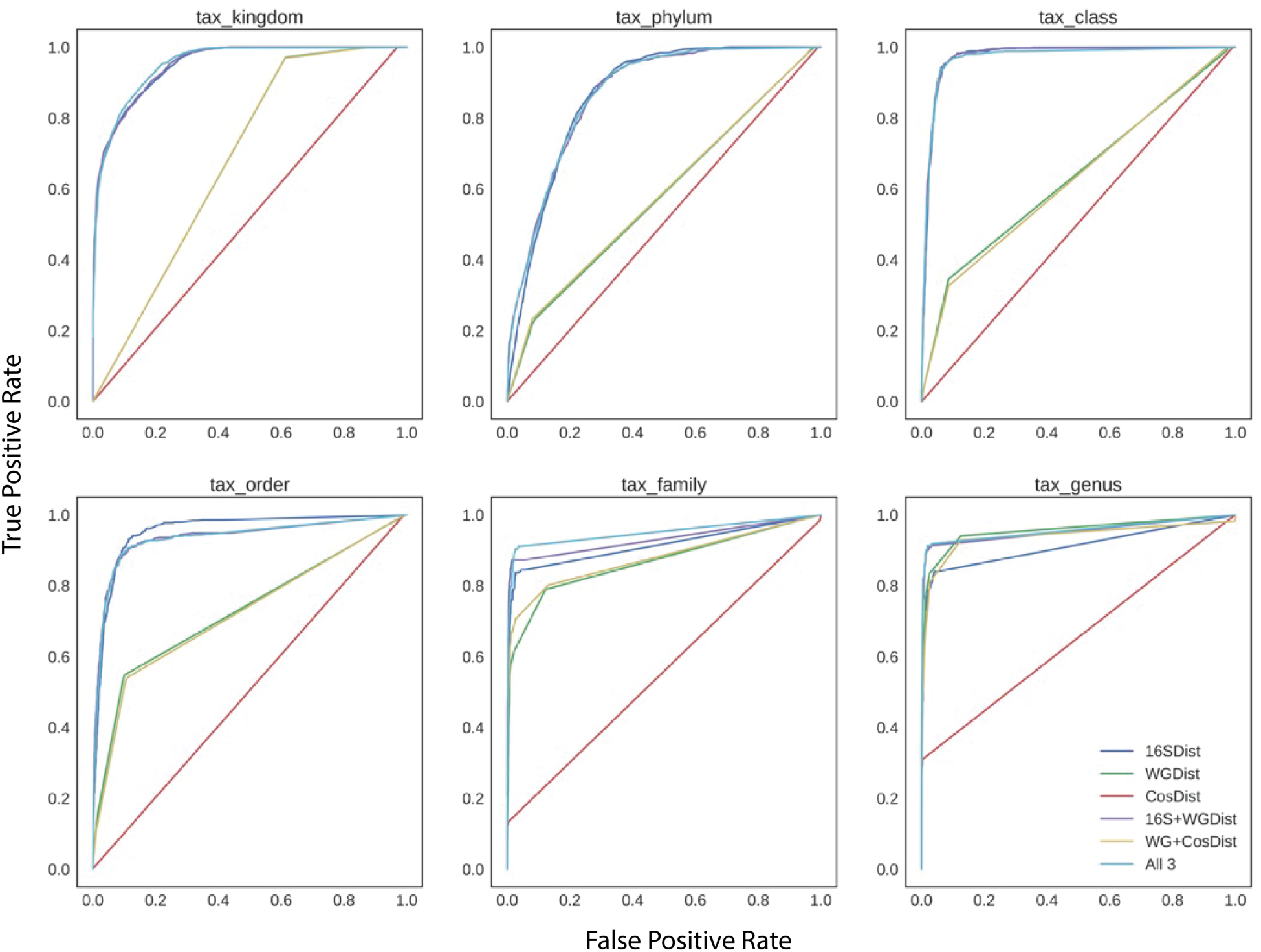
The individual ROC curves for the random forest model trained on each taxonomic distance per pair (0 = pair not in the same taxonomic group, 1 = pair in the same taxonomic group). Each curve represents the data input given (16SDist: 16S MASH distance, WGDist: whole genome distance, CosDist: synteny distance).

**Supplementary Figure 5.**
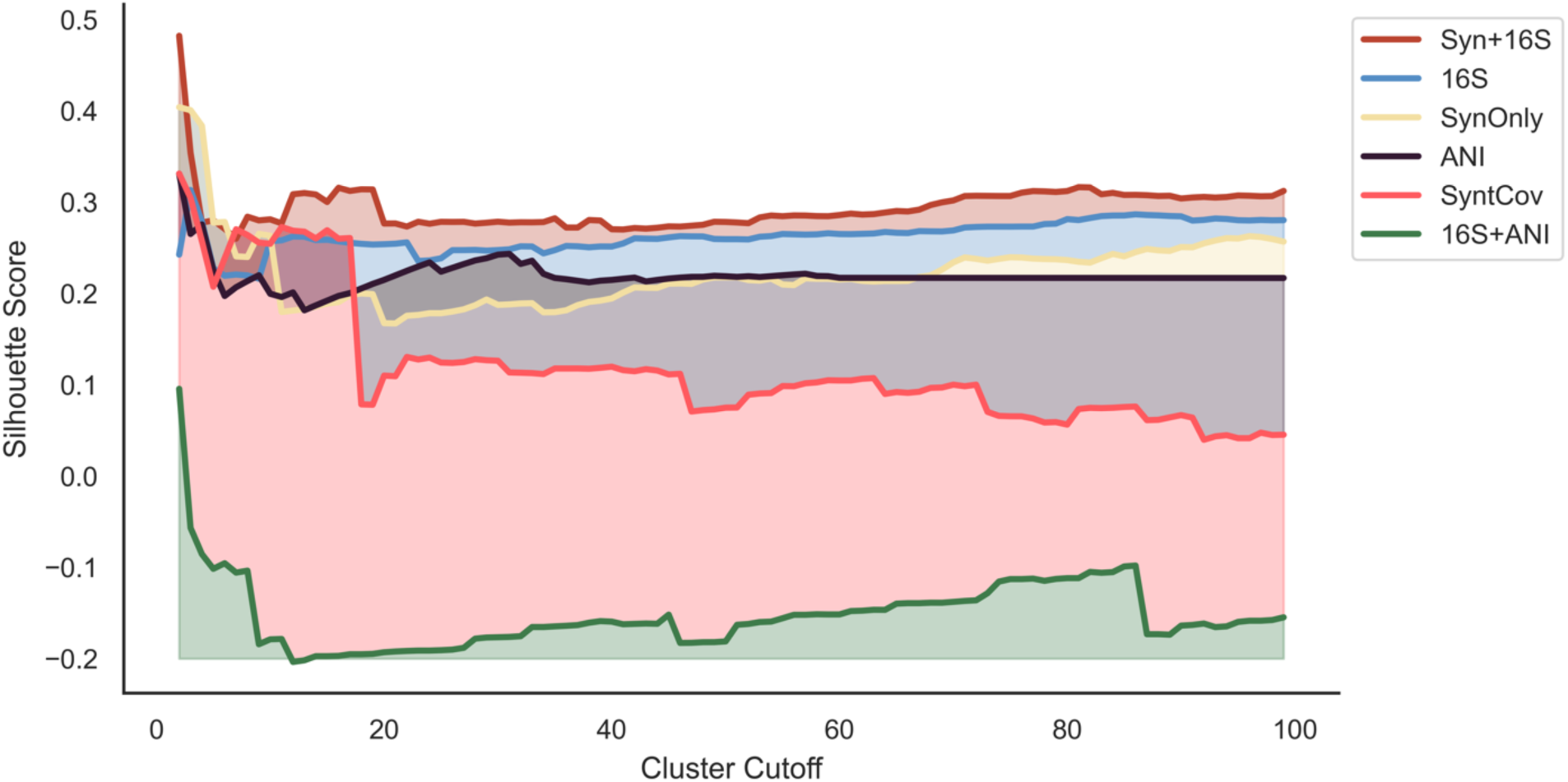
Silhouette score comparison for complete-linkage hierarchical clustering. Cohort names indicate as follows - Syn+16S: the novel synteny metric added to 16S; 16S: the original 16S unchanged matrix; SynOnly: only the original synteny matrix, not added to 16S; ANI: average nucleotide identity matrix; SyntCov: the synteny coverage matrix added to 16S; and 16S+ANI: ANI added to 16S using the same covariance metric tactic.

**Supplementary Figure 6.**
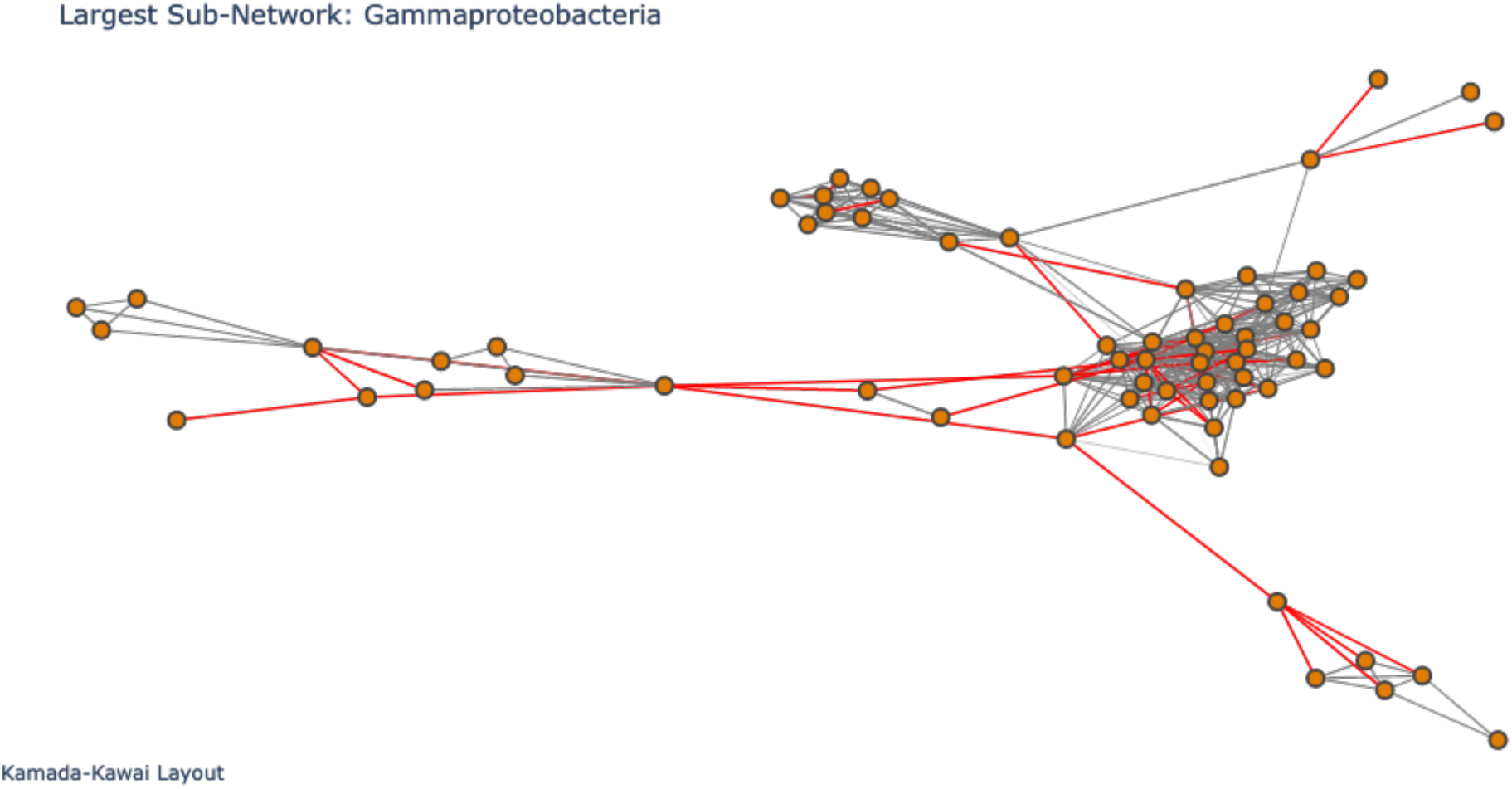
Largest sub-network from synteny network (Figure 2). Edges indicate a similarity value present based on synteny. Edge color is determined by weight (synteny similarity). Red edges have weight > 0.9 and edges below are grey. All nodes in the network are part of the class of bacteria, *Gammaproteobacteria*.

**Supplementary Figure 7.**
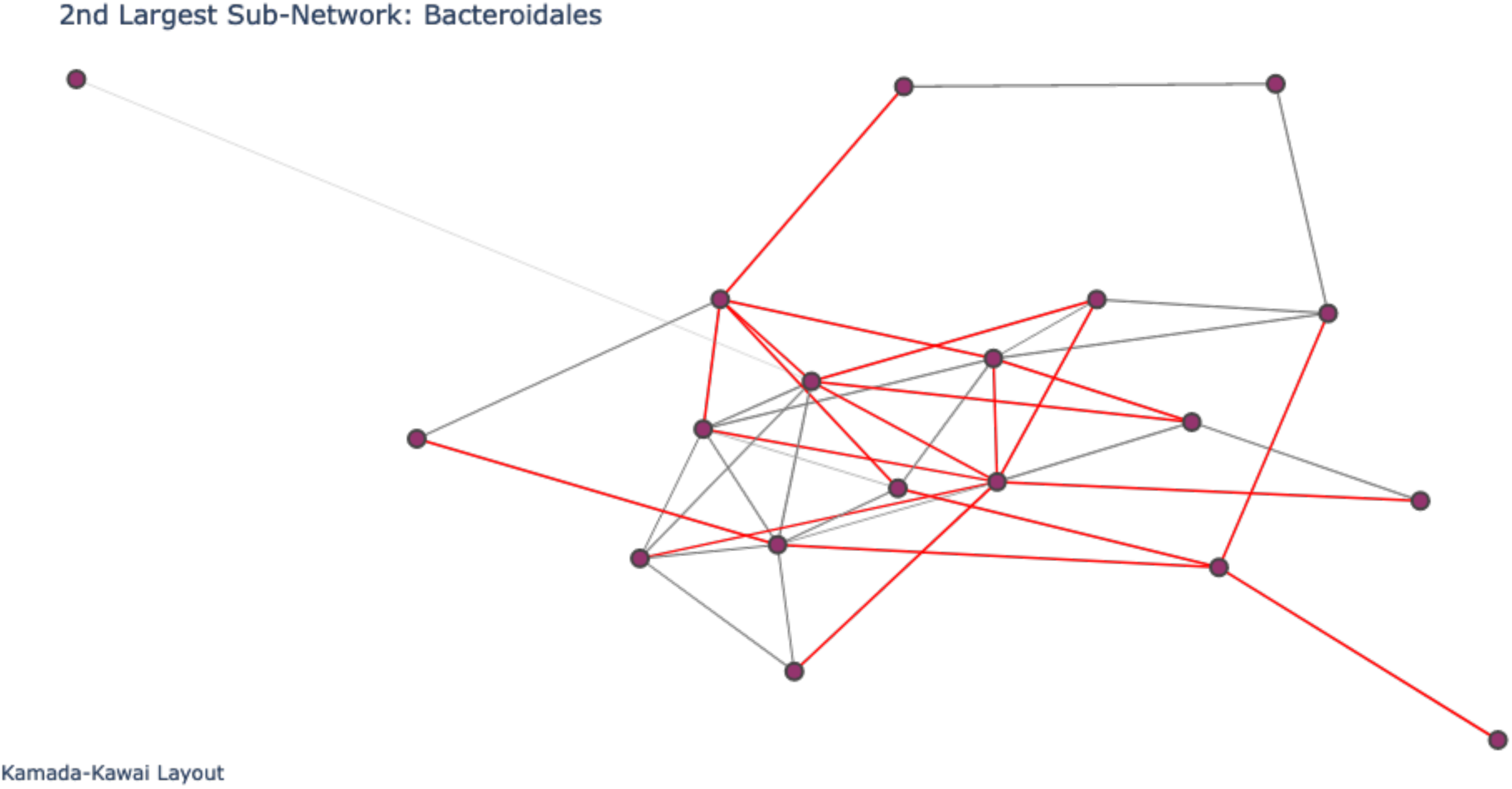
Second largest sub-network from synteny network (Figure 2). Edges indicate a similarity value present based on synteny. Edge color is determined by weight (synteny similarity). Red edges have weight > 0.9 and edges below are grey. All nodes in the network are part of the order of bacteria, *Bacteroidales*.

**Supplementary Figure 8.**
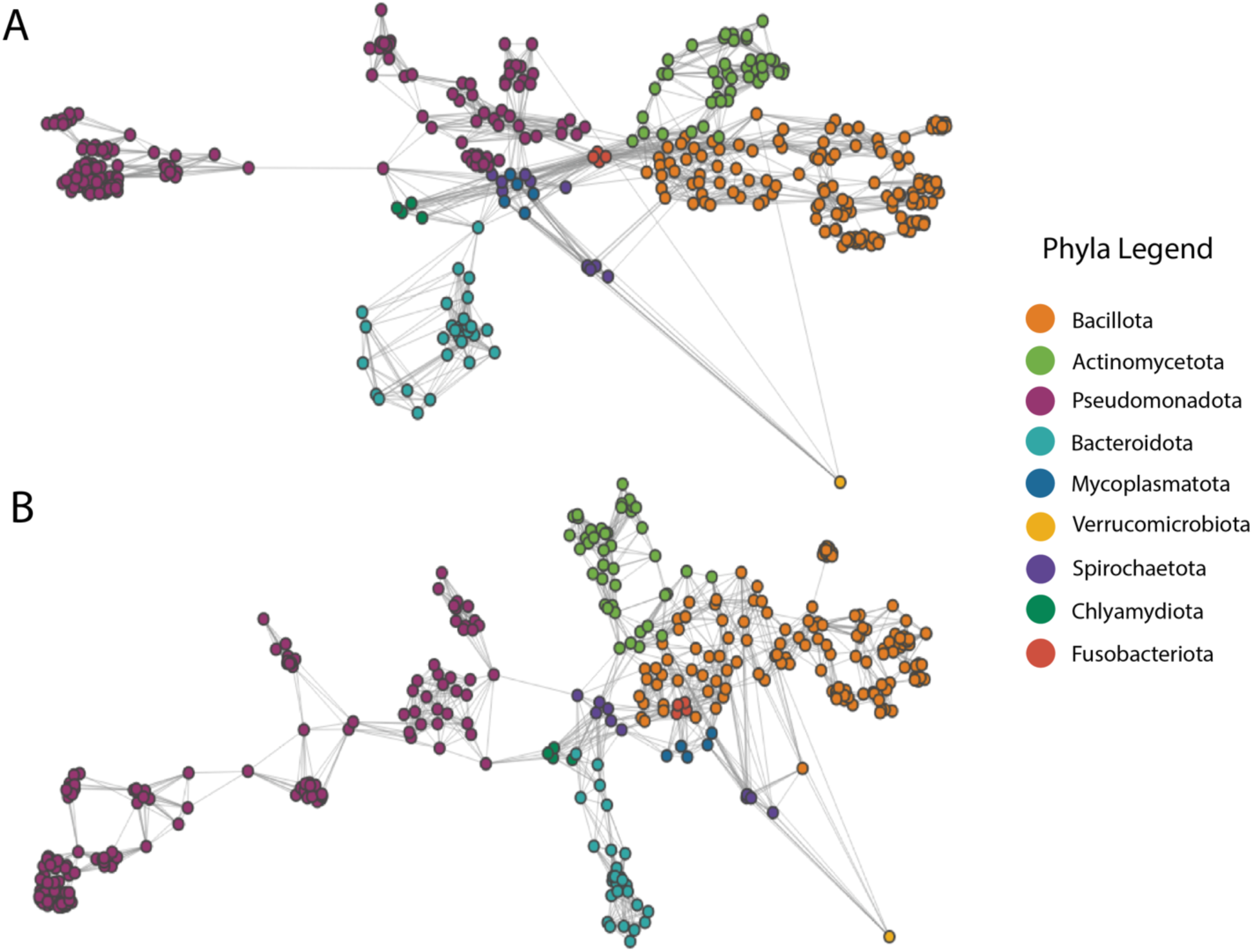
Visualizations of k-nearest neighbor graphs for A) 16S rRNA MASH distance data and B) synteny-scaled 16S data. Nodes represent taxa species, which are labeled by taxonomical phyla.

**Supplementary Figure 9.**
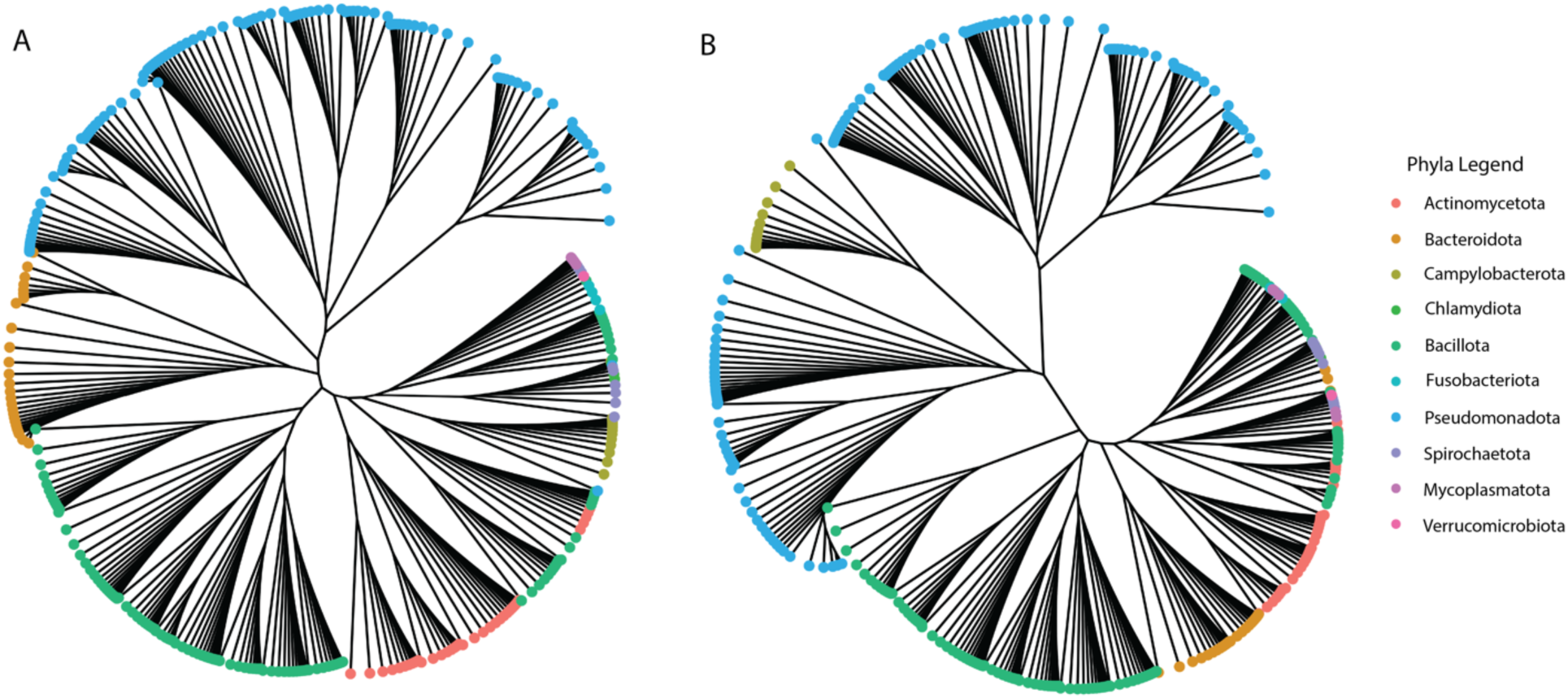
Dendrograms based on Girvan-Newman hierarchical clustering, with A) the 16S rRNA distance data and B) the synteny-scaled 16S data. Leaves of the tree branches are labeled with the taxonomical phyla of the species genome.

**Supplementary Table 1.** Clustering quality descriptive statististics. Silhouette scores used for hierarchical clustering and modularity used from KNN graphs. Averaged across cluster cutoff range 2-100 for clusters and 1-25 for KNN. Cohort definitions can be found in Methods section.

| **METRIC** | **COHORT** | **MEAN** | **SD** | **MEDIAN** |
| --- | --- | --- | --- | --- |
| **SILHOUETTE** | Thr | 0.437 | 0.028 | 0.438 |
|  | Rem | 0.365 | 0.036 | 0.352 |
|  | Syn+16S | 0.294 | 0.025 | 0.286 |
|  | 16S | 0.263 | 0.017 | 0.263 |
|  | ANI | 0.219 | 0.017 | 0.217 |
| **MODULARITY** | Thr | 0.769 | 0.080 | 0.759 |
|  | Rem | 0.163 | 0.162 | 0.093 |
|  | Syn+16S | 0.694 | 0.102 | 0.673 |
|  | 16S | 0.702 | 0.096 | 0.665 |
|  | ANI | 0.162 | 0.135 | 0.107 |
| **SILHOUETTE** | ARG | 0.443 | 0.025 | 0.451 |
|  | MET | 0.445 | 0.015 | 0.440 |
|  | VIR | 0.388 | 0.015 | 0.389 |
|  | MGE | 0.381 | 0.027 | 0.374 |
|  | SYN | 0.294 | 0.025 | 0.286 |
| **MODULARITY** | ARG | 0.635 | 0.133 | 0.576 |
|  | MET | 0.633 | 0.128 | 0.598 |
|  | VIR | 0.653 | 0.111 | 0.612 |
|  | MGE | 0.648 | 0.118 | 0.597 |
|  | SYN | 0.694 | 0.102 | 0.673 |

**Supplementary Table 2.** Sparsity values of each distance matrix provided. Cohort definitions remain the same as all tables.

| **COHORT** | **SPARSITY** |
| --- | --- |
| **THR** | 0.993 |
| **REM** | 0.993 |
| **SYN+16S** | 0.002 |
| **16S** | 0.586 |
| **ANI** | 0.993 |
| **SYNTENY ONLY** | 0.989 |
| **ARG** | 0.934 |
| **MGE** | 0.858 |
| **MET** | 0.926 |
| **VIR** | 0.907 |

**Supplementary Table 3.** Descriptive statistics of comparison metrics used for each cohort against ANI (average nucleotide identity). Rand scores used for clustering and weighted jaccard for KNN graphs.

| **METRIC** | **COHORT** | **MEAN** | **SD** | **MEDIAN** |
| --- | --- | --- | --- | --- |
| **RAND SCORE** | 16S-ANI | 0.559 | 0.137 | 0.591 |
|  | Syn-ANI | 0.567 | 0.132 | 0.595 |
|  | Rem-ANI | 0.859 | 0.047 | 0.841 |
|  | Thr-ANI | 0.525 | 0.114 | 0.546 |
| **WEIGHTED JACCARD** | 16S-ANI | 0.707 | 0.034 | 0.696 |
|  | Syn-ANI | 0.664 | 0.057 | 0.666 |
|  | Rem-ANI | 0.708 | 0.047 | 0.685 |
|  | Thr-ANI | 0.580 | 0.075 | 0.563 |
| **RAND SCORE** | ARG-ANI | 0.538 | 0.117 | 0.564 |
|  | MGE-ANI | 0.551 | 0.127 | 0.581 |
|  | MET-ANI | 0.532 | 0.115 | 0.556 |
|  | VIR-ANI | 0.527 | 0.113 | 0.551 |
| **WEIGHTED JACCARD** | ARG-ANI | 0.645 | 0.017 | 0.642 |
|  | MGE-ANI | 0.660 | 0.018 | 0.660 |
|  | MET-ANI | 0.685 | 0.016 | 0.687 |
|  | VIR-ANI | 0.660 | 0.016 | 0.658 |

## Notes

### Competing Interest Statement

The authors have declared no competing interest.

